# A Universal Probe Set for Targeted Sequencing of 353 Nuclear Genes from Any Flowering Plant Designed Using k-medoids Clustering

**DOI:** 10.1101/361618

**Authors:** Matthew G. Johnson, Lisa Pokorny, Steven Dodsworth, Laura R. Botigue, Robyn S. Cowan, Alison Devault, Wolf L. Eiserhardt, Niroshini Epitawalage, Félix Forest, Jan T. Kim, James H. Leebens-Mack, Ilia J. Leitch, Olivier Maurin, Douglas E. Soltis, Pamela S. Soltis, Gane Ka-Shu Wong, William J. Baker, Norman J. Wickett

## Abstract

Sequencing of target-enriched libraries is an efficient and cost-effective method for obtaining DNA sequence data from hundreds of nuclear loci for phylogeny reconstruction. Much of the cost associated with developing targeted sequencing approaches is preliminary data needed for identifying orthologous loci for probe design. In plants, identifying orthologous loci has proven difficult due to a large number of whole-genome duplication events, especially in the angiosperms (flowering plants). We used multiple sequence alignments from over 600 angiosperms for 353 putatively single-copy protein-coding genes to design a set of targeted sequencing probes for phylogenetic studies of any angiosperm lineage. To maximize the phylogenetic potential of the probes while minimizing the cost of production, we introduce a k-medoids clustering approach to identify the minimum number of sequences necessary to represent each coding sequence in the final probe set. Using this method, five to 15 representative sequences were selected per orthologous locus, representing the sequence diversity of angiosperms more efficiently than if probes were designed using available sequenced genomes alone. To test our approximately 80,000 probes, we hybridized libraries from 42 species spanning all higher-order lineages of angiosperms, with a focus on taxa not present in the sequence alignments used to design the probes. Out of a possible 353 coding sequences, we recovered an average of 283 per species and at least 100 in all species. Differences among taxa in sequence recovery could not be explained by relatedness to the representative taxa selected for probe design, suggesting that there is no phylogenetic bias in the probe set. Our probe set, which targeted 260 kbp of coding sequence, achieved a median recovery of 137 kbp per taxon in coding regions, a maximum recovery of 250 kbp, and an additional median of 212 kbp per taxon in flanking non-coding regions across all species. These results suggest that the Angiosperms353 probe set described here is effective for any group of flowering plants and would be useful for phylogenetic studies from the species level to higher-order lineages, including all angiosperms.

## Plant Phylogenetics and Reduced Representation Sequencing

Progress in molecular phylogenetics has frequently been a struggle between the availability of genetic markers and the suitability of those markers for the specific systematic study. This is especially true in plants, where numerous gene and genome duplication events (Blanc and Wolfe 2004; Barker et al. 2009; Jiao et al. 2011; Amborella Genome Project et al. 2013) have made identification of universally orthologous genes difficult. As a result, phylogenetic inference in plants has frequently relied on plastid markers, using either single genes such as *rps4* or *rbc*L (Palmer et al. 1988; Chase et al. 1993; Soltis et al. 1993; Olmstead and Sweere 1994), entire plastid exomes (Ruhfel et al. 2014; Gitzendanner et al. 2018; Medina et al. 2018) or the full plastid genome sequence (Carbonell-Caballero et al. 2015; Bernhardt et al. 2017). Although sequence homology can easily be determined for the plastid genome, thus being universally applicable across plants, it is often considered to represent a single phylogenetic history (though there are exceptions, see (Sullivan et al. 2017) that may be incongruent with species trees inferred from nuclear genes (Hudson 1992; Maddison 1997). In cases where nuclear data have been applied to plant phylogenetics, it has generally relied upon relatively few loci (e.g. ITS, reviewed in (Alvarez and Wendel 2003); low copy loci reviewed in (Zimmer and Wen 2012). However, studies of both empirical and simulated data demonstrate that phylogenetic inference is most accurate when conducted with tens to hundreds of nuclear loci because historical processes such as deep coalescence can be modeled (Degnan and Rosenberg 2006; McCormack et al. 2009; Smith et al. 2015).

Several reduced-representation sequencing methods have been developed to sample hundreds of nuclear loci for plant phylogenetic studies (reviewed in (McKain et al. 2018). These methods allow users to reap the benefits of high-throughput sequencing, yielding data sets of tractable scale for phylogenetics without the bioinformatic challenges and costs associated with, for example, whole genome sequencing. Aside from cost considerations, the decision of which method to use depends on the taxonomic breadth of the study as well as the availability of existing genomic resources. For example, restriction site-associated sequencing methods (RAD-Seq or GBS) are an efficient and cost-effective way to generate single nucleotide polymorphisms (SNPs) for lower-level phylogenetic studies without having to rely on existing genome or transcriptome sequences (Eaton et al. 2016). However, the markers generated by this method are exclusive to the lineage for which they were developed and may introduce a number of biases (reviewed in (Andrews et al. 2016)). Another reduced representation method that has been employed for plant phylogenetics is transcriptome sequencing (Wickett et al. 2014; Yang et al. 2015; Zeng et al. 2017; Walker et al. 2018). Despite reduced costs compared to whole genome sequencing, and increased efficiency in both library construction and sequencing, transcriptomes are not the most cost-effective and reproducible source of data for phylogenetics. Assembled transcripts include members of multi-copy gene families and other genes that may not be phylogenetically informative, therefore sequencing cost and effort is not optimized for the reconstruction of species phylogenies. Furthermore, the same genes may not be expressed in the same tissues for all targeted taxa, reducing reproducibility and increasing the amount of missing data. Finally, generating transcriptomic data sets requires fresh tissues and is not feasible for extremely rare, extinct, or ancient samples, where only herbarium specimens may be available.

Plastid data have long met the criteria of reproducibility and cost-effectiveness necessary for phylogenetics in non-model plants. In addition, the widespread use of specific plastid loci for plant phylogenetics has facilitated data reuse in analyses of expanded data sets and higher-order relationships (e.g., (Chase et al. 1993). A set of similarly consistent, overlapping, cost-effective nuclear loci would enable plant systematists to produce data appropriate for lineage-specific investigations while generating data that would benefit the plant systematics community more broadly. By populating sequence databases with the same set of putatively orthologous loci, taxon-rich analyses that mine *all* publicly available sequences (Hinchliff and Smith 2014; Eiserhardt et al. 2018; Smith and Brown 2018) will be subject to lower levels of missing data. The question remains, then, as to which reduced-representation method best addresses the goal of generating an accessible and combinable set of loci for plant phylogenetics.

## Target Enrichment, Considerations, and Objectives

Among genome-scale methods developed to date, the sequencing of target-enriched genomic libraries has emerged as a cost-effective method for obtaining large data sets for phylogenetics from diverse sources, particularly for methods that rely on reconstructing gene trees (Faircloth et al. 2012; Lemmon et al. 2012; Mandel et al. 2014; Weitemier et al. 2014). In this method, high-throughput sequencing libraries are enriched for regions of interest, such as ultra-conserved elements or protein-coding genes, using 80–120-mer DNA or RNA probes (sometimes called baits) that hybridize to library inserts. Like the design of primer regions for Sanger sequencing, the selection of loci and design of probe sequences requires a careful balance: the challenge of selecting genomic regions variable enough to infer phylogenies while remaining conserved enough to ensure sequence recovery. As discussed above, the unique challenges of identifying universal phylogenetically informative loci for plants (Kadlec et al. 2017) have likely contributed to the relatively small number of targeted sequencing probe sets applicable across a broad phylogenetic spectrum (e.g. all vascular plants or all flowering plants) compared to the availability of such tools in animals (Faircloth et al. 2012; Lemmon et al. 2012; Prum et al. 2015; Faircloth 2017).

A set of orthologous single-copy genes identified from 25 angiosperm genomes was recently used to develop probes for target enrichment across angiosperms (Léveillé-Bourret et al. 2018). Although there are about 75 well-assembled and annotated angiosperm nuclear genomes publicly available (e.g., Phytozome version 12.1, www.phytozome.org), the phylogenetic distribution of genomic resources is uneven and groups that are not closely related to economically important flowering plants are poorly sampled. To maximize the potential for successful hybridization to probe sequences from a set of phylogenetically diverse species, it is critical to include as much phylogenetic breadth as possible when designing probes. For example, projects such as the Plant and Fungal Trees of Life (www.paftol.org) —which aims to capture the diversity of all angiosperm genera (13,164 genera; (Christenhusz and Byng 2016))— or the Genealogy of Flagellate Plants (http://flagellateplants.group.ufl.edu/) —which aims to sequence all flagellate plant species (ca. 30,000 species of bryophytes, lycophytes, ferns, and gymnosperms; (Villarreal et al. 2014; Forest et al. 2018))— require a universal set of markers that represent an even distribution of diversity. To maximize the likelihood of successful hybridization in any angiosperm lineage, the design of a universal probe set for angiosperms should make use of the most phylogenetically diverse set of gene sequences available.

In contrast to available genomic resources, transcriptome sequences generated by the 1000 Plant Transcriptomes Initiative (OneKP or 1KP) provide a more even phylogenetic distribution (Matasci et al. 2014) and include sequences for over 830 flowering plant taxa (data available at onekp.com/public_data.html). Transcriptome sequences have been successfully used to develop probe sets for targeting nuclear protein-coding genes in several plant groups (Chamala et al. 2015; Landis et al. 2015, 2017; Gardner et al. 2016; Heyduk et al. 2016; Crowl et al. 2017; García et al. 2017; Stubbs et al. 2018). Although intron-exon boundaries are not known when probes are designed exclusively from transcriptomes in non-model organisms, this does not prevent efficient sequence recovery (Heyduk et al. 2016). The design of probes to capture coding sequences commonly results in the capture of non-coding sequence flanking the exons, (i.e. the “splash-zone”; (Weitemier et al. 2014); Fig. 1). This protocol is useful for narrow-scale phylogenetic analysis (Heyduk et al. 2016; Johnson et al. 2016) and represents an advantage over transcriptomes due to the consistent and reproducible recovery of more variable non-coding regions.

**Figure 1.**
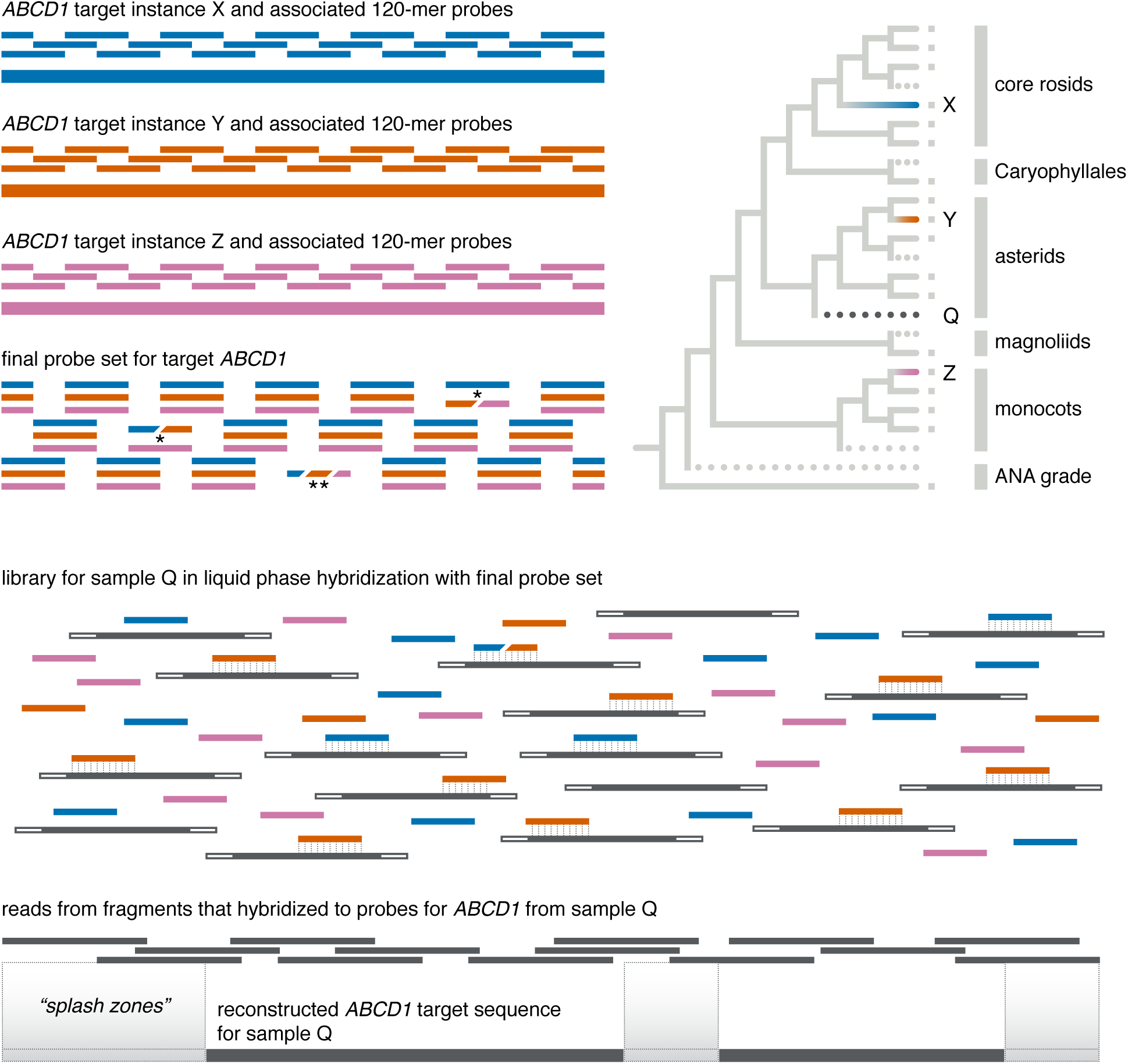
Overview of Probe Design and Phylogenetic Considerations. Given a hypothetical gene *ABCD1*, the goal of probe design is to include a sufficient diversity of 120-mers (probes) such that any angiosperm *ABCD1* sequence can be recovered by hybridization. If a number of *ABCD1* sequences are known, represented by solid branches and tips with a small gray square in the phylogeny, the minimum number of representatives of those sequences should be selected that maximize the chances of recovering *ABCD1* from any “unknown” sample (dotted lines in the phylogeny). If sequences X, Y, and Z are selected, 120-mer probes are designed, here with 2X tiling, across the entire length of the sequence. The final probe set includes all unique 120-mers; asterisks represent cases in which individual 120-mers are identical from two (^∗^) or all three (^∗∗^) of the representative sequences X, Y, and Z. In these cases only one or two 120-mers, rather than three, would be necessary for that region of the gene in the final probe set. While this is possible in probe design, we did not encounter any such cases in the Angiosperms353 probe set. For a particular “unknown” sample, here represented by the dark gray dotted line and denoted as sample Q, a sequencing library consisting of size-selected inserts, adapters, and indexes, is hybridized to the final probe set and the resulting sequence reads can be reconstructed to extract both the coding region and flanking non-coding (“splash zone”) regions. In this simplified example, the final probe set represents only gene *ABCD1* but the Angiosperms353 final probe set includes probes tiled across 353 genes.

Designing probes for target enrichment of nuclear protein-coding genes involves the identification of three pieces of information: (1) the locus itself or **target** gene, ideally one that exists as a single copy in all species belonging to the lineage under investigation; (2) the minimum number of **target instances** such that the target genes from any input sample are sufficiently related to ensure hybridization, and (3) **probe** sequences designed from all selected target instances, tiled across the length of each instance (Figure 1). For example, the target genes and all target instances could be full length (whenever possible) coding sequences (CDS) of genes inferred to be orthologous across a lineage, while the probes are RNA sequences of known length (usually ~120-mers) that hybridize to genomic DNA, usually sequencing library inserts. Multiple probes are tiled across the target instances to ensure that the entire target is recovered. If a substantial phylogenetic breadth of input libraries is to be used for hybridization, multiple target instances must be present in the probe set to maximize the chance that inserts will hybridize with at least one probe from a specific region of the target, since the chance of hybridization is proportional to sequence similarity between the input library and probes. A “universal” set of probes, therefore, would first require identifying a suitable number of sequences to “universally represent” a target sequence for a large lineage (Figure 1).

Flowering plants are estimated to have arisen sometime in the Early Cretaceous (Bell et al. 2010; Herendeen et al. 2017; Barba-Montoya et al. 2018) and comprise approximately 369,400 extant species (Kew 2016). Therefore, for any given target sequence, probes designed from several instances of that target sequence would be required for the successful hybridization of inserts from taxa that are significantly divergent from any representative in the probe set. For example, if a hypothetical gene *ABCD1* is a target, probes may need to be designed from a monocot *ABCD1*, a rosid *ABCD1*, and an asterid *ABCD1*, at a minimum, to ensure that *any* angiosperm *ABCD1* sequence would successfully hybridize with at least one set of probes that span the entire length of *ABCD1* (Figure 1). Because increasing the number of probes increases the cost of a target enrichment kit, producing a cost-effective kit to enrich phylogenetically informative exons from any angiosperm requires minimizing the number of instances of each target sequence while maximizing the phylogenetic depth of hybridization.

We report the design of a set of target enrichment probes that efficiently capture hundreds of putatively orthologous gene regions (targets) from any angiosperm species. Our two main objectives were to: (1) develop an approach to choose the minimum number of target instances needed to successfully recover the targets from any flowering plant, and (2) generate probes from those target instances and use empirical data to demonstrate that there is no phylogenetic bias in the probe set. We introduce here a novel application of a k-medoids clustering algorithm for selecting target instances. The target instances chosen via this method allowed for a probe design using fewer than 80,000 probes to capture 353 protein-coding genes. We carried out an initial test of the probe sequences on 42 species spanning 30 angiosperm orders. We find high rates of recovery for both the targeted coding sequences and flanking intron regions, suggesting that the probe set will be a cost-efficient and universally accessible tool for flowering plant phylogenetics at both deep and shallow scales.

## Probe Design

### Target Identification

We started with a set of 410 protein-coding loci developed for phylogenetic analysis by the OneKP initiative (One Thousand Plant Transcriptomes Initiative, *In Review(Matasci et al. 2014).* All available green plant (Viridiplantae) genomes were clustered into homologous gene families using OrthoFinder (Emms and Kelly 2015). For each family, a Hidden Markov Model was generated from a multiple sequence alignment, and transcriptome sequences from over 1400 green plant species were added to the gene families using hmmsearch implemented in HMMER (hmmer.org). The 410 alignments identified as low-copy contained orthologous transcripts from over 1100 green plants and predicted coding sequences from 52 plant genomes. We removed all non-angiosperm sequences from the OneKP alignments and trimmed all gap-only sites from the alignments. The 410 nucleotide alignments contained between 12 and 655 angiosperm sequences and varied in length from 105 to 3498 bp.

### Selection of Target Instances

To minimize the number of probe sequences needed to reliably recover sequences from all angiosperms, we reduced the OneKP alignments by selecting the minimum possible number of target instances. Our goal was to select instances such that 95% of all angiosperm sequences in the alignment were no more than 30% diverged from any target instance, a threshold that has been demonstrated as the practical limit of target enrichment in other plant groups (Li, Johnson et al., in review). First, we calculated a dissimilarity matrix (p-distance) using all pairwise non-gap characters in the angiosperm-only sequence alignments. We employed a k-medoids clustering algorithm (Bauckhage 2015) to partition the sequences into groups, centered around a set number of sequences (the medoids). The k-medoid method attempts to minimize the within-group distance between the medoid and other sequences in the group. We chose this method over the related k-means clustering method because the k-medoid approach would identify a single real sequence at the center of each cluster, rather than a hypothetical sequence at the centroid of a cluster generated by k-means. We explored how varying the number of medoids (k) affected the percentage of angiosperm sequences that could be representative of the whole alignment (at a maximum 30% sequence divergence). For each gene, we tested values of k between 5 and 15, repeating each analysis up to 100 times. To evaluate the k-medoids method, we calculated the sequence divergence between all angiosperm transcript sequences and the selected medoid sequences and compared this to the sequence divergence from manually selected target instances chosen from publicly available genome sequences (Figure 2). The scripts used to select k-medoid alignments are available at http://github.com/mossmatters/Angiosperms353.

**Figure 2.**
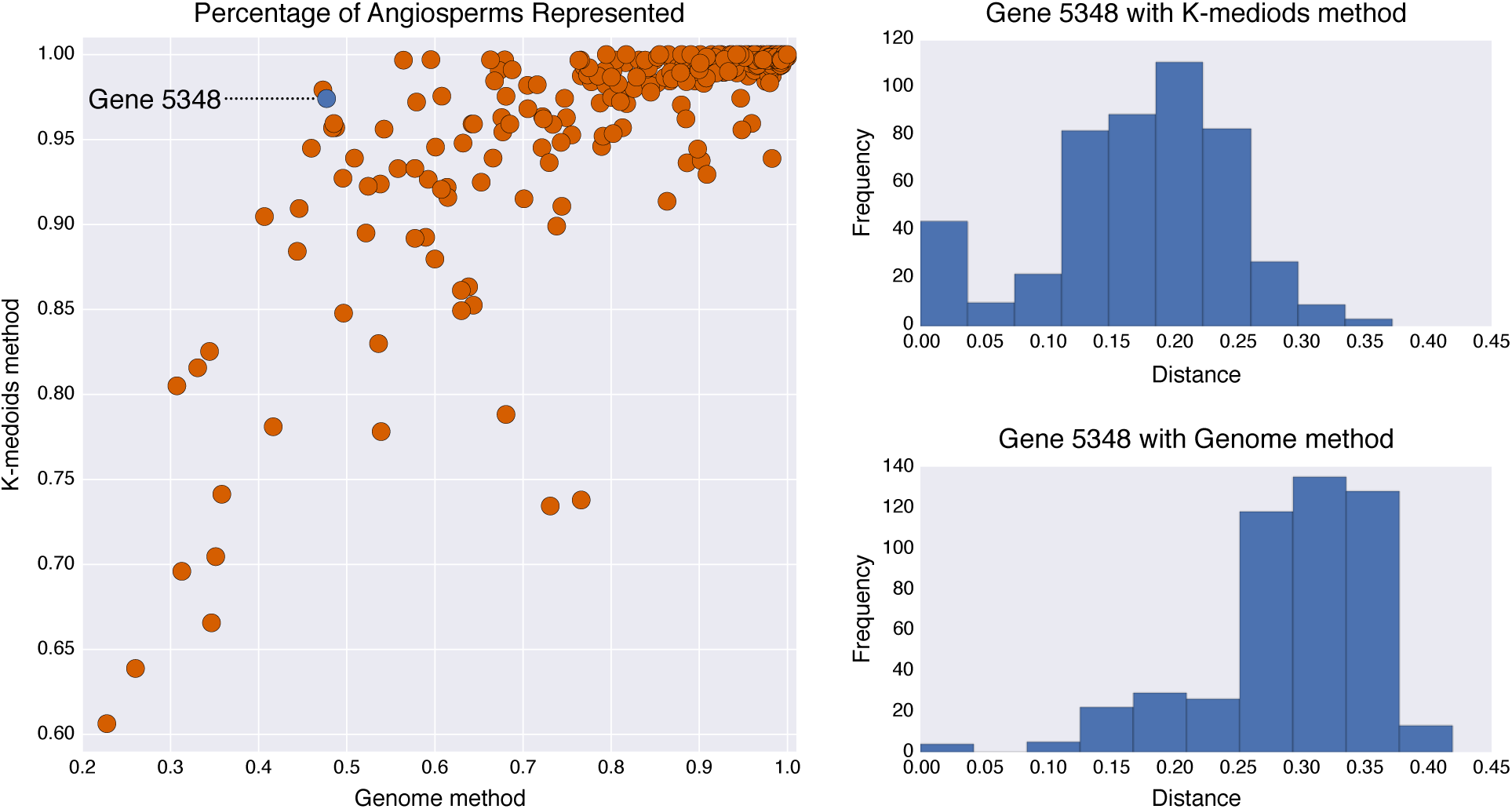
Comparison Between the k-medoids Method of Selecting Representative Sequences with Using the Closest Available Angiosperm Genome. A) Each point is one gene, and its position indicates the percentage of angiosperm transcripts (from OneKP) that fall within 30% sequence divergence of a representative sequence. Only genes where the k-medoids could represent 95% or more angiosperms were selected for probe design. Note the x-axis and y-axis ranges are not identical. Arrow indicates gene 5348, which is highlighted in the other panels. B) Distribution of distances between each angiosperm sequence in 1KP and the nearest k-medoid for gene 5348. C) Distribution of distances between each angiosperm sequence in 1KP and the nearest published genome sequence for gene 5348.

### Final Probe Set

Using the k-medoids method, we identified 353 genes (targets) for which 95% of angiosperm sequences could be represented by 15 or fewer target instances (Figure 2). A sequence was represented if it was within 30% sequence divergence of one of the target instances. For each gene, if they were not already in the set of representative sequences chosen by the k-medoid method, we added target instances from the genomes of *Arabidopsis thaliana*, *Oryza sativa*, and *Amborella trichopoda.* Sequences from these three genomes were added to ensure a well-annotated gene model was present for each gene spanning deep divergences in the angiosperm phylogeny. Probe sequences were designed from multiple sequence alignments of all selected target instances for each gene. Across all 353 targets, there were 4781 sequences (target instances) used for probe design (all alignments are available in the Dryad repository). Probe sequences were padded to a minimum length of 120 bases, and any regions with 1-10 Ns were replaced with Ts (to facilitate probe design across short stretches of ambiguity). Synthesis of 120-mer RNA probes with 3x tiling on each orthologous sequence was carried out by Arbor Biosciences (formerly MYcroarray, Ann Arbor, Michigan, USA, Catalog #3081XX). Each probe sequence was compared to seven angiosperm reference genomes (*Amborella trichopoda*, *Aquilegia coerulea*, *Brachypodium distachyon*, *Populus trichocarpa*, *Prunus persica*, *Solanum tuberosum*, and *Vitis vinifera)* to check for specificity; any probe sequence with multiple hits in more than one genome was removed from the probe set. The final probe set contains 75,151 120-mer probes. Probe sequences are publicly available under a CC-BY-SA license at github.com/mossmatters/Angiosperms353

The goal of our probe design was to maximize the usability of the probes across angiosperms by ensuring that 95% of known angiosperm sequences had less than 30% sequence divergence from any target instance for each gene. We also wanted to minimize the number of target instances needed for each gene to reduce the cost for users of the probes. To validate whether k-medoids method was an improvement over selecting target instances using existing angiosperm genome sequences only, we calculated the pairwise sequence divergence between each OneKP transcript and its nearest ortholog from an angiosperm genome. For each gene, we compared the percentage of angiosperm sequences within 30% divergence of a k-medoid to the percentage within 30% divergence of an angiosperm genome sequence. Using reference genome sequences alone was insufficient and frequently resulted in a large number of angiosperm sequences falling beyond the 30% sequence divergence threshold (Figure 2). Using the k-medoids method, we were able to successfully identify a different number of target instances for each gene, based on individual sequence divergence profiles across angiosperms. The number of required medoid sequences ranged from six (in 64 genes) to 15 (in 125 genes) with an average of 11.1 medoid sequences (median 13 sequences). Of the 410 original angiosperm alignments, medoids fitting our criteria could not be identified for 57 genes, leaving 353 loci in the final probe design. Using the mean length of target instances from each gene, the total length of coding sequence targeted was 260,802 bp.

## Testing the Probe Set

### Sampling Strategy and Data Generation

To test the efficiency of target capture across angiosperms, we tested the probes using 42 “input” taxa that were not included in the OneKP transcriptome data (Table 1), increasing the likelihood that no input insert would be identical to any probe sequence. However, *Amborella trichopoda* was also included as an input taxon as a control of recovery efficiency by using a taxon that was included in probe design. Our sampling scheme was designed to test whether target recovery is determined by relatedness to the species used to design the probes (i.e. to ensure that the probes are not phylogenetically biased). We used taxonomic rank from APG IV (The Angiosperm Phylogeny Group 2016) as a proxy for phylogenetic distance. For each input species, its relationships to the OneKP taxa (pool of potential probe sequences) fell into four categories, which we refer to throughout as **Input Categories**:

1. The input taxon belongs to an order that was not included in OneKP (7 taxa)
2. The input taxon belongs to an order that was included in OneKP, but not to a family, genus, or species that was included in OneKP (12 taxa)
3. The input taxon belongs to an order and family that were included in OneKP, but not to a genus or species that was included in OneKP (12 taxa)
4. The input taxon belongs to an order, family, and genus included in OneKP, but not to a species that was included in OneKP (11 taxa).

**Table 1.**
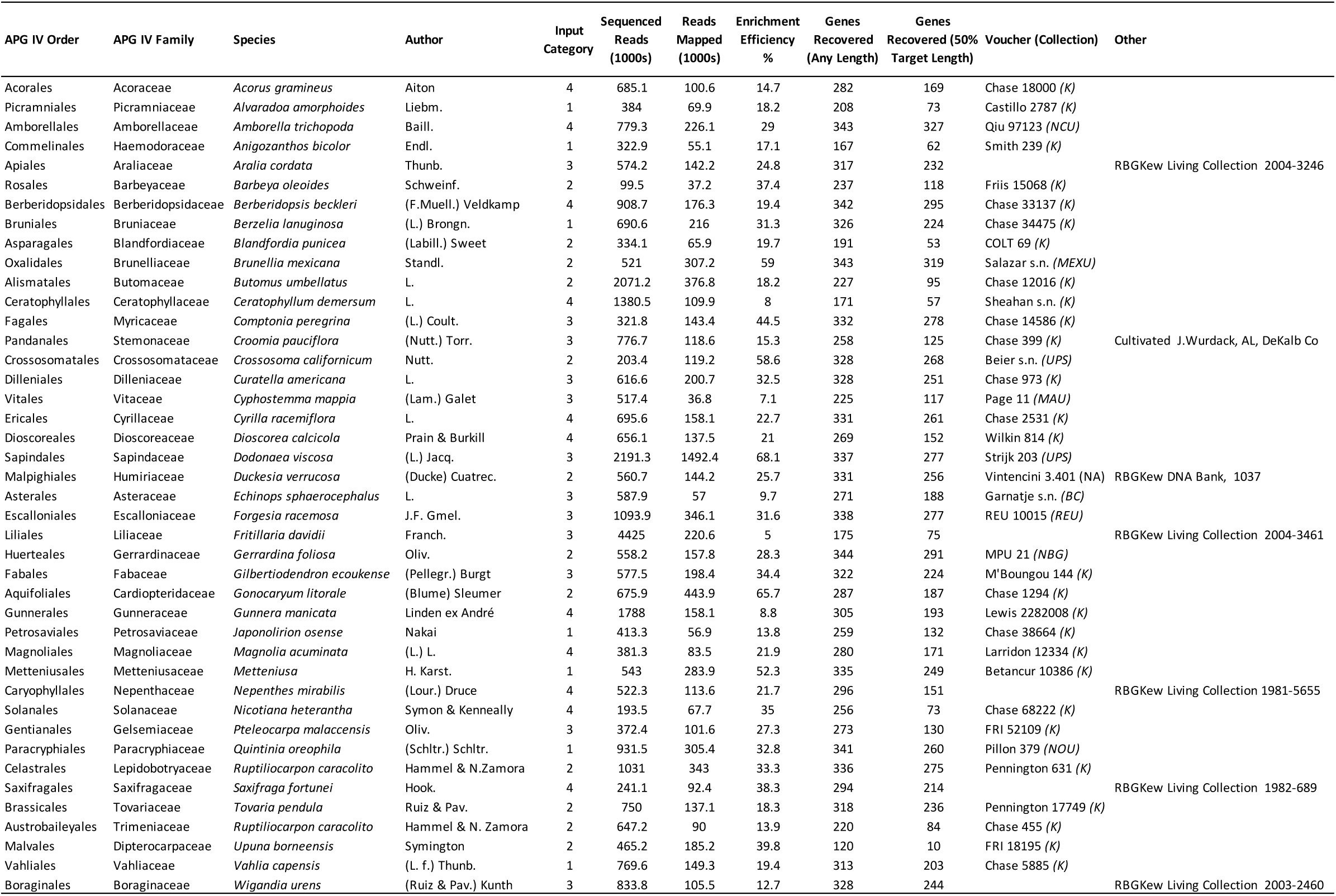
Voucher Information, Targeted Sequencing Efficiency, and Gene Recovery for 42 Angiosperms.

The input taxa span 41 of the 64 angiosperm orders recognized in APG IV. If the probe set is phylogenetically biased, we would expect to see any metrics of hybridization efficiency improve from Input Category 1 to Input Category 4. Conversely, if hybridization success is more dependent on stochastic effects, we would expect to see no relationship between Input Category and hybridization efficiency.

DNA extractions were tailored to tissue provenance: the Qiagen DNeasy Plant Mini Kit was used for silica-dried materials following the manufacturer’s protocol (Qiagen, Valencia, California, USA), while a modified CTAB protocol (Doyle and Doyle 1987) was the choice for material sampled from herbarium specimens. We also relied on existing DNA extractions from the DNA bank of the Royal Botanic Gardens, Kew, obtained using a standard CTAB-chloroform, ethanol precipitation and washing stages, followed by density gradient cleaning and dialysis. All extractions were run on a 1.5× agarose gel to visually assess average fragment size and quantified using a Qubit^®^ 3.0 Fluorometer (Life Technologies, Carlsbad, California, USA). Samples demonstrated (by gel) to have fragment sizes above the desired 350 bp (typically those obtained from silica-dried tissue and most DNA bank aliquots) were sonicated using a Covaris M220 Focused-ultrasonicator™ with Covaris microTUBES AFA Fiber Pre-Slit Snap-Cap (Covaris, Woburn, Massachusetts, USA) following the manufacturer’s program for ~350-bp insert sizes. Dual-indexed libraries for Illumina^®^ sequencing were prepared using the DNA NEBNext^®^ Ultra™ II Library Prep Kit at half the recommended volume, with Dual Index Primers Set 1, NEBNext^®^ Multiplex Oligos for Illumina^®^ (New England BioLabs, Ipswich, Massachusetts, USA). All resulting libraries were checked for quality with an Agilent Technologies 4200 TapeStation System using the High Sensitivity D1000 ScreenTape (Agilent Technologies, Santa Clara, California, USA) and quantified with the Qubit^®^ 3.0 Fluorometer. Equimolar 1 μg pools were enriched using our custom-designed probe kit (Arbor Biosciences myBaits^®^ Target Capture Kit, “Angiosperms 353 v1”, Catalog #3081XX) following the manufacturer’s protocol (ver. 3, available at http://www.arborbiosci.com/mybaits-manual). Multiple libraries were pooled in a single hybridization reaction. Hybridizations were carried out at 65°C for 28–32 hours in a Hybex^®^ Microsample Incubator with red Chill-out™ Liquid Wax (Bio-Rad, Hercules, California, USA) to prevent evaporation. Enriched products were amplified with KAPA HiFi 2X HotStart ReadyMix PCR Kit (Roche, Basel, Switzerland) for 10 cycles. PCR products were cleaned using the QIAquick PCR purification kit (Qiagen). Final products were quantified with a Qubit^®^ 3.0 Fluorometer and run on an Agilent Technologies 4200 TapeStation System to assess quality and average fragment size. Multiple enriched library pools were multiplexed and sequenced on an Illumina MiSeq with v2 (300-cycles) and v3 (600-cycles) chemistry (Illumina, San Diego, California, USA) at the Royal Botanic Gardens, Kew.

Sequencing reads were trimmed using Trimmomatic (Bolger et al. 2014) to remove reads with a quality score below 20 and reads that had any 4-bp window below 20, retaining reads with at least 50 bp (LEADING: 20 TRAILING: 20 SLIDING WINDOW:4:20 MINLEN:50). Only read pairs where both reads passed these filters were retained. Recovered target sequences were assembled using HybPiper version 1.3 (Johnson et al. 2016) using a target file available at http://github.com/mossmatters/Angiosperms353. Reads were mapped to de-gapped medoid sequences using BWA (Li and Durbin 2009), each gene was assembled de-novo using SPAdes (Bankevich et al. 2012), and coding sequences were extracted using Exonerate (Slater and Birney 2005). Non-coding sequences (i.e. introns) flanking the coding sequences were recovered using the script *intronerate.py* available with HybPiper. Statistical analysis of sequence recovery was conducted in R (Version 3.4.2, R Core Development Team, 2017). Sequence reads have been deposited in the NCBI Sequence Read Archive (SRP151601); recovered gene sequences, alignments, and distance matrices are available from the Dryad Digital Repository (http://dx.doi.org/10.5061/dryad.[NNNN]). Code for statistical analysis is available at http://github.com/mossmatters/Angiosperms353.

### Target Enrichment Results

We assessed the probe set using two main statistics: (1) target enrichment, measured as the percentage of reads successfully mapping to a target instance and (2) gene recovery rate, measured as the percentage of targeted genes recovered by HybPiper. Target enrichment ranged from 5% (*Fritillaria davidii*) to 68% (*Dodonaea viscosa*), with an average of 27.5% (median 24.8%). However, poor enrichment efficiency did not always predict poor gene recovery rate. Coding sequences were recovered from between 120 and 344 loci (average 283); more than 300 genes were recovered from 21 of our 42 samples (Figure 3). The median number of genes for which the length of coding sequence recovered was at least 50% of the target length (the average length of target instances for each gene) was 118 (range 10-327, Table 1). There was no relationship between Input Category (taxonomic relatedness to samples in 1KP) and the number of sequenced reads (F = 0.70, DF = 3, p > 0.5), the number of mapped reads (F = 0.93, DF = 3, p > 0.4), the percentage of reads on target (F = 1.4, DF = 3, p > 0.25), or the number of genes with recovered sequences (F = 0.22, DF = 3, p > 0.8). The lack of relationship between enrichment success and Input Category suggests that there is no phylogenetic bias in the probe set.

**Figure 3.**
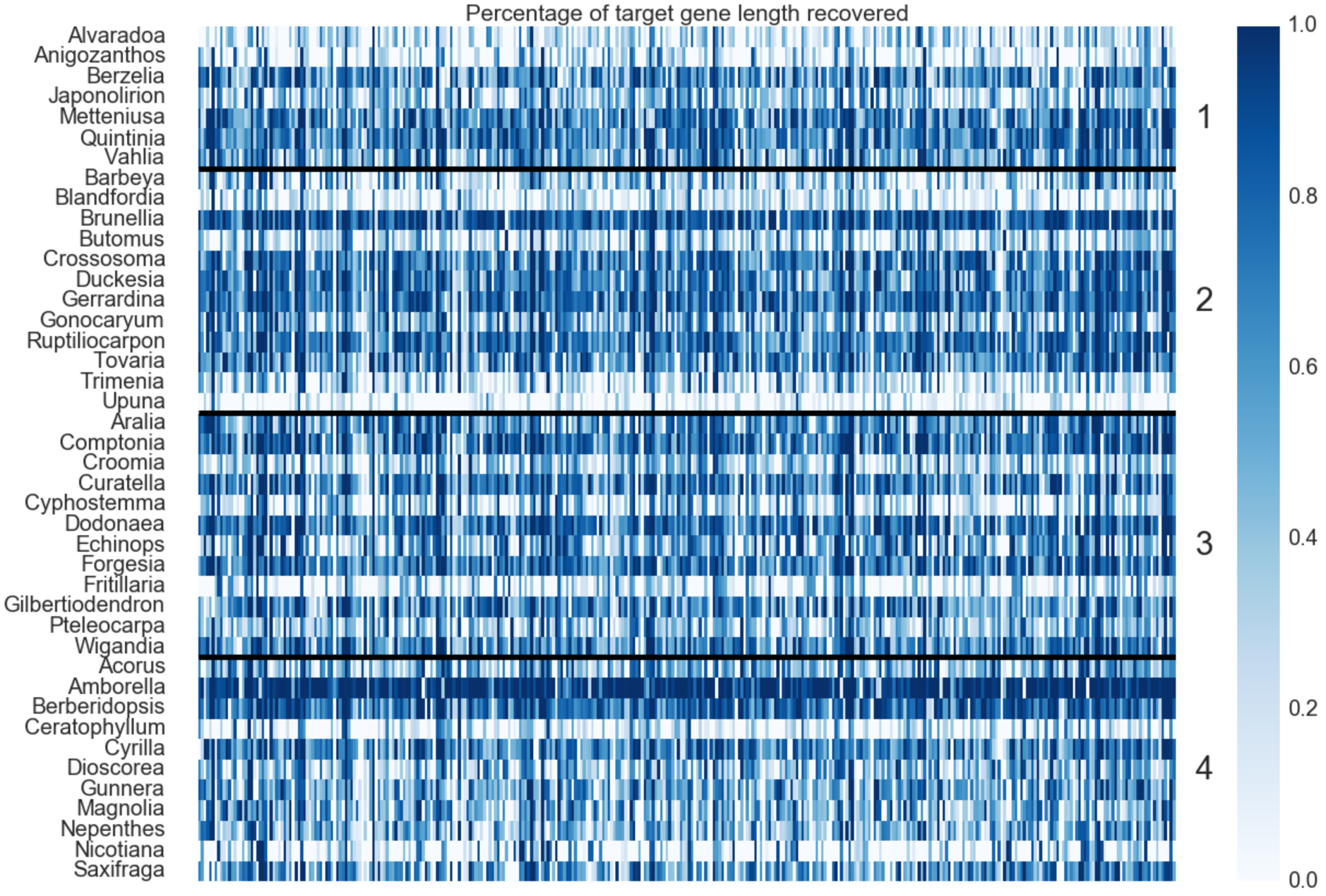
Heatmap of Gene Recovery Efficiency. Each row is one sample, and each column is one gene. Colors indicate the percentage of the target length (calculated by the mean length of all k-medoid transcripts for each gene) recovered. Numbers indicate the Input Category (see main text).

There was no significant linear (F = 0.91, DF = 40, p > 0.3) or log-transformed (F = 0.14, DF = 40, p > 0.7) relationship between the number of genes recovered and the number of sequenced reads. However, there was a significant relationship between the number of genes recovered and the log-transformed number of mapped reads (F = 10.1, DF = 40, p < 0.005). With fewer than 200,000 mapped reads on target, there was roughly a linear relationship between mapped reads and genes recovered, reaching a plateau of roughly 300 genes recovered (Supplemental Figure 1). Therefore, it appears that stochasticity related to the enrichment and sequencing process (e.g., library complexity, pooling strategy), not Input Category, is the most critical factor in determining successful target enrichment (Table 1). This indicates that an input sample with low enrichment efficiency could be “rescued” by increased sequencing effort or by increasing the complexity of the genomic library (for example, by increasing DNA input into library preparation). If the probes exhibited phylogenetic bias, increasing the number of reads or increasing library complexity would not overcome the absence of sufficiently homologous probes, a case that is not reflected in our results.

### Suitability for Lower-Order Analysis

Although our sampling focused on the applicability of the probes across the breadth of angiosperm diversity, we also considered whether the same target genes could be used at narrower phylogenetic scales. Within-genus data are already available from OneKP for the Angiosperms353 target genes from *Oenothera* (Myrtales, Onagraceae, 19 taxa), *Linum* (Malpighiales, Linaceae, 9 taxa), *Portulaca* (Caryophyllales, Portulacaceae, 11 taxa), and *Neurachne* (Poales, Poaceae, 6 taxa). For each genus, we reduced the OneKP alignments to retain only the sequences from that genus and calculated the number of variable characters. In each genus, several dozen variable characters could be found for a large number of genes, with total numbers of variable characters ranging from 30,479 to 109,068 within genera (Table 2, Supplemental Table 1). Overall, the percentage of protein-coding sites that were variable ranged from 7.8% in *Neurachne* to 27.2% in *Linum*, while *Oenothera* and *Portulaca* were intermediate at 10.0% and 13.5%, respectively (Table 2).

**Table 2:** Summary of Parsimony-informative Sites in Multiple Sequence Alignments of Transcripts Sequenced in OneKP for Four Angiosperm Genera at 353 Loci.

| Family | Genus | Taxa | Total Sites | Variable Sites |
| --- | --- | --- | --- | --- |
| Onagraceae | *Oenothera* | 19 | 386152 | 38618 |
| Portulacaceae | *Portulaca* | 9 | 386427 | 52217 |
| Poaceae | *Neurachne* | 6 | 392432 | 30479 |
| Linaceae | *Linum* | 11 | 400001 | 109068 |

Sequence capture of coding regions from genomic DNA will also recover non-coding regions (often introns) flanking the exons. This “splash zone” is less constrained by purifying selection and is likely to be useful for narrow-scale phylogenetics and population genetics (Weitemier et al. 2014; Johnson et al. 2016). Although our sampling scheme to test phylogenetic bias in the probes did not allow for direct observation of phylogenetic information within genera, we did recover a large fraction of non-coding flanking sequence (Figure 4). The median amount of non-coding sequence recovered with at least 8× depth of coverage was 216,816 bp, and fluctuated from 32,233 in *Upuna* (Category 2) to 664,222 bp in *Amborella* (Category 4). Input Category had no significant effect on the length of non-coding regions recovered (df = 38, F = 0.16, p > 0.9). The combination of variable sites within coding regions among congeners and the significant recovery of flanking non-coding regions suggests that this probe set will be valuable for reconstructing both relationships at both shallow and deep phylogenetic scales.

**Figure 4.**
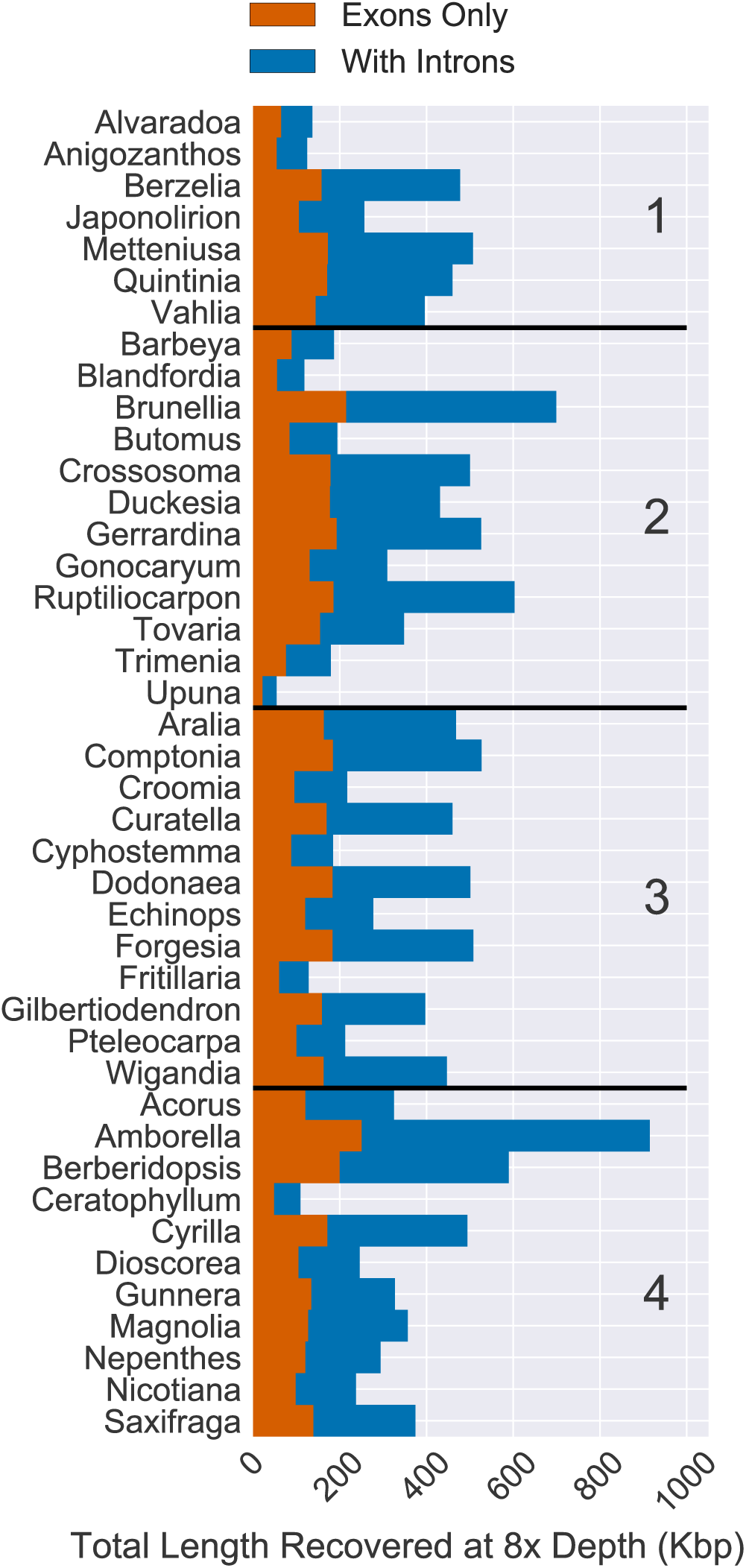
Total Length of Sequence Recovery for Both Coding and Non-coding Regions Across 353 Loci for 42 Angiosperm Species. Reads were mapped back to either coding sequence (yellow) or coding sequence plus flanking non-coding (i.e. intron) sequence (purple). Only positions with at least 8× depth were counted. The total length of coding sequence targeted was 260,802 bp. The median recovery of coding sequence was 137,046 bp and the median amount of non-coding sequence recovered was 216,816 bp (with at least 8x depth of coverage).

## Availability and Ongoing Development

Probe sequences for the “Angiosperms 353 v1” (Arbor Biosciences Catalog #3081XX) angiosperm-wide targeted sequencing kit are publicly available at github.com/mossmatters/Angiosperms353. We expect that updates to the kit will be made to improve target enrichment efficiency across angiosperms, and these changes will be tracked as new versions on github. For example, some additional consideration may be made for groups of angiosperms with high phylogenetic distance to other groups (e.g., *Ceratophyllum* and *Gunnera*), which may necessitate additional orthologous probe sequences. The sequences of target instances used are also freely available at the same site and will be similarly updated to help improve sequence recovery. For example, as the use of the kit increases the availability of sequences in poorly represented groups, these sequences may be added to the target sequence file used to reconstruct sequences in software such as HybPiper.

## Conclusions

The creation of a universal set of target enrichment probes for angiosperms requires a phylogenetically diverse set of existing genomic or transcriptomic resources and a method that can identify the minimum set of representatives from which probes can be designed. Here, we have shown that the k-medoids method with a diverse set of existing transcriptome sequences can identify a suitable number of representatives for target enrichment across a wide phylogenetic breadth. By designing probes based on these representatives and testing them empirically, we have demonstrated the potential of our probe set to be a universal DNA sequencing resource across all angiosperms. While sequence recovery efficiency varied among samples, these differences are not driven by relatedness between the input taxon and any probe sequence, indicating that there is no phylogenetic bias to sequence recovery across angiosperms using these probes. Instead, the recovery of gene sequences was impacted more by the number of reads mapped per library, suggesting that critical samples with low gene recovery could be improved by further sequencing, or by a more careful evaluation of library complexity. The data presented here indicate that these probes will be useful for phylogenetic analysis at both deep and shallow scales, particularly given the recovery of flanking non-coding regions. Additional applications of the universal probe set could extend to inferring population level genetic structure, or to the next generation of DNA barcoding (Hollingsworth et al. 2016). Given that hybridization-based methods of library enrichment using the probes described here should be successful for most, if not all, angiosperms, there is significant potential for the generation of large, combinable data sets for future analyses.

## Acknowledgements

This work was supported by funding from the Texas Tech University College of Arts and Sciences to MGJ, grants from the National Science Foundation (DEB-1239992, DEB-1342873) to NJW, and by grants from the Calleva Foundation, the Garfield Weston Foundation, and the Sackler Trust to the Royal Botanic Gardens, Kew.

## Supplemental Material

Supplemental Table 1. Sequence length, number of taxa, and parsimony informative sites in alignments of transcriptome sequences at 353 genes for four angiosperm genera.

Supplemental Figure 1. Relationship between reads mapping to the target genes and the number of loci recovered for 42 angiosperm species. There is a general linear increase in the number of genes recovered below 100,000 mapped reads, above which there are diminishing returns for additional sequencing.

